# Genomic Distance-based Rapid Uncovering of Microbial Population Structures (GRUMPS): a reference free genomic data cleaning methodology

**DOI:** 10.1101/2022.12.19.521123

**Authors:** Kaleb Z. Abram, Zulema Udaondo, Michael S. Robeson, Se-Ran Jun

**Affiliations:** Department of Biomedical Informatics, University of Arkansas for Medical Sciences, Little Rock, Arkansas, USA

**Keywords:** Population structure, phylogenomics, Mash, comparative genomics, taxonomy, Average Nucleotide Identity, data cleaning, k-means clustering, machine learning, big data

## Abstract

Accurate datasets are crucial for rigorous large-scale sequence-based analyses such as those performed in phylogenomics and pangenomics. As the volume of available sequence data grows and the quality of these sequences varies, there is a pressing need for reliable methods to swiftly identify and eliminate low-quality and misidentified genomes from datasets prior to analysis. Here we introduce a robust, controlled, computationally efficient method for deriving species-level population structures of bacterial species, regardless of the dataset size. Additionally, our pipeline can classify genomes into their respective species at the genus level. By leveraging this methodology, researchers can rapidly clean datasets encompassing entire bacterial species and examine the sub-species population structures within the provided genomes. These cleaned datasets can subsequently undergo further refinement using a variety of methods to yield sequence sets with varying levels of diversity that faithfully represent entire species. Increasing the efficiency and accuracy of curation of species-level datasets not only enhances the reliability of downstream analyses, but also facilitates a deeper understanding of bacterial population dynamics and evolution.

## Introduction

With sequencing technologies becoming increasingly affordable and accessible worldwide the amount of publicly available whole genome sequences is skyrocketing. Heavily sequenced species such as *Escherichia coli*, now have tens of thousands of sequences publicly available which can, nevertheless, overwhelm local computational resources when utilizing a variety of analytical methods, such as alignment-based analytical methodologies^1^. While there are over 300,000 bacterial assembled genomes available in public repositories such as GenBank (as of September 2024), large microbial datasets may exhibit inherent sampling and sequencing biases that are virtually impossible to avoid^2,3^. These biases stem from the prioritization of sequencing efforts towards clinically relevant strains, important for human health, and model organisms, influenced by funding availability^3^. Furthermore, biases may arise from preferential sampling of easily cultivable microbes, geographical considerations, host associations, methodological differences, and historical research trends, collectively influencing the representation and quality of sequences in repositories^4^.

Analyses performed using biased datasets can lead to conclusions about the genomic traits of bacterial species that fail to capture the true diversity of the sequenced species. Similarly, the inclusion of poor-quality genome sequences in analyses can yield incorrect conclusions or mask biologically meaningful results^5^. Conversely, disregarding a large portion of available sequences at the species level to mitigate these biases, would ultimately constrain the insights obtainable from comprehensive analysis. This is particularly crucial for comparative analyses reliant on gene presence thresholds, such as comparative genomics, pangenome studies, taxonomy, and core genome multi-locus sequence typing analyses. Ensuring access to unbiased curated datasets will enable more accurate and informative exploration of microbial genomics^6–10^.

To facilitate the performance of large-scale whole genome sequence (WGS) analyses, computational tools such as FastANI^11^ and Mash^12^ have been developed to enhance efficiency. These tools aim to overcome computational challenges inherent in pairwise comparisons across extensive datasets by rapidly estimating genetic similarity between bacterial genomes. However, their focus lies in speeding up the pairwise comparisons process by utilizing k-mers and MinHash techniques, without directly providing high-quality genomic datasets. Similarly, while various methodologies, metrics, and tools exist to evaluate genome quality (e.g., CheckM^13^, for completeness and contamination, presence/absence of gene markers, N50/NG50, and L50/LG50 ratios)^14^ they lack automation for large dataset cleaning, particularly in the absence of reference genomes. Without robust methodologies addressing this gap, the efficacy of tools such as FastANI^11^ and Mash^12^ will remain constrained by the inclusion of low-quality or mislabeled genomic sequences.

To address these challenges, we have developed GRUMPS (**<u>G</u>**enetic distance based **<u>R</u>**apid **<u>U</u>**ncovering of **<u>M</u>**icrobial **<u>P</u>**opulation **<u>S</u>**tructures), a Python-based program engineered to provide a novel approach to rapidly and reproducibly clean bacterial datasets of any magnitude. GRUMPS represents an enhancement and extension of the methodology recently employed to conduct the largest population structure analyses of *E. coli* and *Pseudomonas aeruginosa* to date^10,15^. Leveraging a diverse array of automated and statistical techniques, GRUMPS systematically removes outlier genomes from bacterial species datasets. The majority of the cleaning modes provided by GRUMPS rely on statistical analyses and unsupervised machine learning algorithms to accurately detect and exclude divergent genomes within the dataset. Additionally, GRUMPS introduces a mechanism for segregating multiple species within datasets containing multiple bacterial species or genus-level datasets. The resulting isolated species can subsequently undergo further refinement using GRUMPS’ species-level cleaning modes, culminating in the generation of high-quality bacterial species datasets. Moreover, GRUMPS facilitates the reconstruction of population structures within the species under investigation through pairwise genome comparisons across the entire dataset. The genetic structure of a microbial population, or population structure, provides the organized distribution of genetic variations among a set of strains from different geographical locations and ecological niches inside the species^16,17^. Thus, the population structure of a set of strains can be leveraged to better understand the biological context of a bacterial species and the unique traits of subgroups within a given species^10,15^. The genetic structure of a microbial population can also be leveraged to unveil existing biases (such as those from sequencing efforts) within a dataset. Thus, the availability of curated datasets at the species level provided by GRUMPS will offer several advantages, including enhanced accuracy and reliability in downstream analyses, improved understanding of bacterial population dynamics, and greater insights into microbial evolution and adaptation.

## RESULTS

### The theory behind GRUMPS

In theory, a genomic dataset of any bacterial species consists of a collection of genome sequences whose sequence similarity distribution potentially approximates to a normal distribution centered around an overall species-level similarity value. This value ranges from 0 (indicating identical genomes) to 1 (representing genomes with no shared characteristics) for Mash distances. Mislabeled or low-quality genomes would deviate markedly from the similarity values exhibited by the members of the species, falling on the right side of a histogram depicting similarity values for the species. Therefore, we hypothesize that statistics-based methods can reliably identify and exclude genomes that fall outside the normal distribution of similarity values within the dataset. This conclusion would lead to an overall improvement in the quality of the dataset without compromising the sequence diversity of the species.

### Leveraging a hypothetical dataset to illustrate the theory

A hypothetical bacterial species dataset was constructed, containing 1,533 members with a normally distributed range of sequence similarities as measured by Mash distances. This uncleaned hypothetical species dataset has an average Mash distance of 0.045419 which falls below the species boundary of 0.05 Mash distance established by Ondov *et al*.. This boundary is approximately equivalent to a DNA-DNA hybridization value of 70% and an ANI value of 95%^12^. Based on the assumptions described in the previous section, the distribution of the Mash distances for this species should be Gaussian and centered slightly below 0.05, reflecting the average Mash distance of the species (Figure 1).

A histogram representation of the Mash distances for this hypothetical bacterial species reveals a bimodal distribution (Figure 1a). The maximum Mash distance in Figure 1a is 0.087769, which exceeds the species boundary of 0.05 Mash distance and would correspond to an ANI value of approximately 91%. Although a bimodal distribution does not explicitly indicate of one or more outlier genomes, the rightmost peak in Figure 1a surpasses the species boundary of 0.05 Mash distance, indicating the presence of one or more groups of genomes outside the species boundary. These outlier values may belong to mislabeled genomes or draft genome assemblies with low sequence quality due to the presence of contaminant sequences from other species, resulting in lower sequence similarity values compared to other members of this hypothetical species^18^.

The clustered heatmap of Mash distances from Figure 1b, is color-coded based on pairwise Mash distances of the genomes from the dataset, with white colors representing Mash distances near the species boundary of 0.05. Regions in the upper left corner of the clustered heatmap, appearing brownish in Figure 1b, correspond to outlier genomes above the species boundary of 0.05 as defined by Mash. As observed in the dendrogram above the heatmap in Figure 1b, these outlier genomes are separated from the large cluster in the heatmap contributing to the bimodal distribution observed in Figure 1a. Both Figures 1a and 1b are based on the uncleaned distance matrix for this hypothetical species, indicating that the dataset contains genome sequences above the species boundary cut-off. Thus, utilizing this dataset containing outlier sequences in downstream WGS-based analyses could lead to erroneous biological conclusions ^18–20^.

The impact of removing the outlier genomes identified in the hypothetical bacterial dataset (n=383) is visualized in Figures 1c and 1d. As depicted in Figure 1c, the rightmost peak of the histogram in Figure 1a disappears, and the average Mash distances shown in Figure 1d decrease considerably, with a maximum distance for the entire hypothetical dataset of 0.0575621 and a lower mean of 0.0384772. The cleaned dataset also reveals a more detailed population structure compared to the uncleaned dataset shown in Figure 1b, attributed to the automatic scaling of the heatmap colors to the new range of Mash distance values in the heatmap.

**Figure 1.**
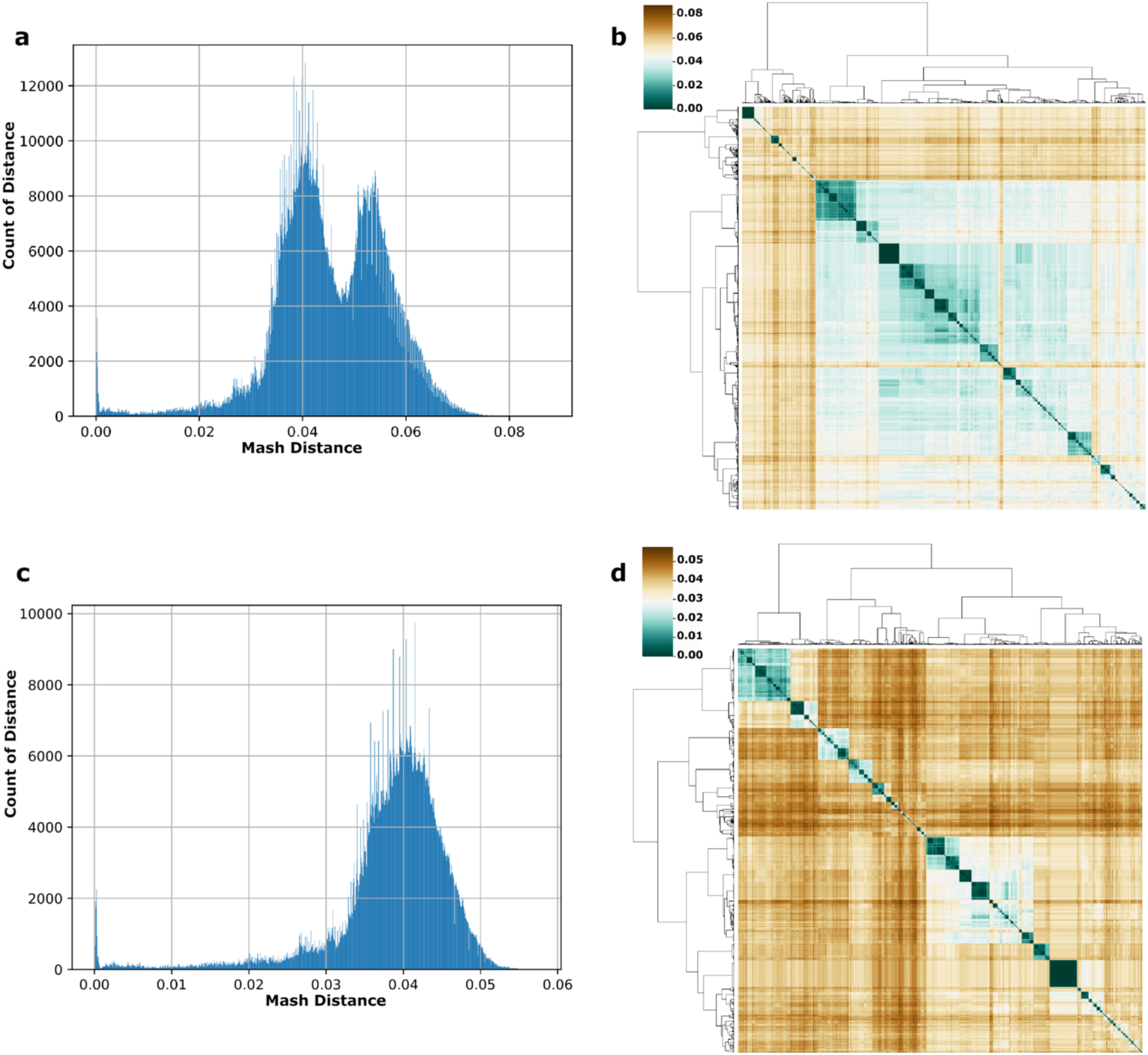
Hypothetical bacterial species before and after cleaning. Panels **a** and **b** depict a dataset consisting of 1,533 genomes belonging to a non-filtered hypothetical bacterial species. **a.** Histogram of Mash values for the entire hypothetical species. The bimodal distribution in panel **a** and the brown region of panel **b**, indicate the presence of low-quality genomes in the dataset. **b.** Clustered heatmap displaying Mash distances for the hypothetical species. Teal colors in both heatmaps represent genomes with higher similarity, while brown colors indicate lower genetic similarity. Relative median similarities are represented in the lightest color. The corresponding Mash distance to color is provided in the key in the upper left-hand corner of each heatmap panel. Panels **c** and **d** illustrate a dataset consisting of 1,170 genomes from the hypothetical bacterial species after removing divergent genomes, which may result from low-sequence quality or mislabeled genomes. **c.** Histogram of Mash values after cleaning and removing outlier genomes from the hypothetical dataset. The disappearance of the bimodal distribution in panel **c** is attributed to the removal of the divergent genomes from the initial dataset. **d.** Heatmap displaying Mash distances for the filtered hypothetical species.

### General Summary of GRUMPS

To apply the methodology used in the hypothetical bacterial dataset and test our hypothesis on real datasets, we developed the GRUMPS (<u>G</u>enetic distance based <u>R</u>apid <u>U</u>ncovering of <u>M</u>icrobial <u>P</u>opulation <u>S</u>tructures) pipeline. GRUMPS is written in Python and designed to be scalable, time efficient, and accessible to the entire scientific community using local computational resources. GRUMPS can be installed via conda, pip, or downloaded from the following GitHub repository: https://github.com/kalebabram/GRUMPS. GRUMPS offers a novel methodology for the rapid and automated cleaning of bacterial genomes datasets. It achieves this by identifying and removing outlier genomes through scalable and reproducible statistical methods, in combination with unsupervised machine learning and user specified cutoffs. GRUMPS is designed to work with distance matrices containing pairwise genome comparisons, with genome IDs (such as GenBank Assembly Accessions) serving as column and row indexes. The values within the distance matrix typically range from 0 to 1, such as those provided by Mash^12^, where a distance of 0 indicates identical genomes and a distance of 1 indicates no shared genetic features.

While GRUMPS was primarily designed to utilize distance matrices generated from Mash output, it also offers helper scripts to assist researchers in transforming their own distance or similarity matrices into a compatible format. Given its design around Mash, as the intended input, GRUMPS defaults to a cutoff of 0.05, determined experimentally as the species boundary for Mash distances^12^. Researchers can adjust this cutoff to suit their specific needs. In instances where Mash distances are not utilized, GRUMPS encourages the use of pairwise comparison methodologies with established species-level cutoffs, such as Average Nucleotide Identity (ANI)^21–23^ or Average Amino Acid Identity (AAI)^24,25^. Furthermore, GRUMPS offers a range of filtering methods to enhance its utility.

### Description of the Different Modes Available in GRUMPS

The “summary” mode serves as the initial step for utilizing GRUMPS. It furnishes various statistical summaries of the input dataset along with a histogram of comparison values, allowing for manual identification of outlier genomes, if desired. If a cutoff value other than the default 0.05 is preferred, we suggest using the sum of the mean and standard deviation output by the “summary” mode as the cutoff value, particularly for datasets with documented smaller genomic variability or significant contamination. This mode can be useful when analyzing species, such as *M, tuberculosis,* whose genomic variability has been documented to be relatively smaller than other species^26^ or datasets with enough contamination that the overall average of the dataset is greater than 0.05. The primary cleaning methods offered by GRUMPS are the “regular” and “strict” modes. Both modes augment the dataset with a small number of controlled artificial outliers, set slightly above the cutoff, to enhance subsequent clustering. After adding these outliers, both modes utilize k-means clustering (k=2) to generate two clusters: one with inlier genomes and the other containing outlier genomes and artificial outliers. The “strict” mode additionally removes any genome with a mean distance greater than the sum of the mean and three times the standard deviation. An optional cleaning step can also be applied. The “sigma” mode employs the three-sigma rule to remove genomes exhibiting divergent behavior relative to the dataset’s distribution of comparisons (both left and right sides). This mode is automatically applied after the “regular”, “strict”, “target”, and “remover” modes. The “target” mode eliminates genomes from the dataset that do not meet the set cutoff distance to specified reference genomes. The “remover” mode is used to eliminate specified genomes from the input dataset, such as known outliers. The “clique” mode partitions datasets with multiple species into smaller uncleaned species-level datasets and unconnected genomes. It trims edges greater than the cutoff from the graph-based dataset, returning subgraphs for cleaning with the “regular” or “strict” modes. For datasets with insufficient members for clustering in “regular” or “strict” modes after using “clique”, the “small” cleaning mode removes outlier genomes based on their average distance. Figure 2a provides an overview of the GRUMPS workflow, with helper scripts available for alternate workflows. Figure 2b summarizes the steps of each cleaning method. A more detailed description of these modes and their specific methodology is provided in the Supplementary Material section.

**Figure 2.**
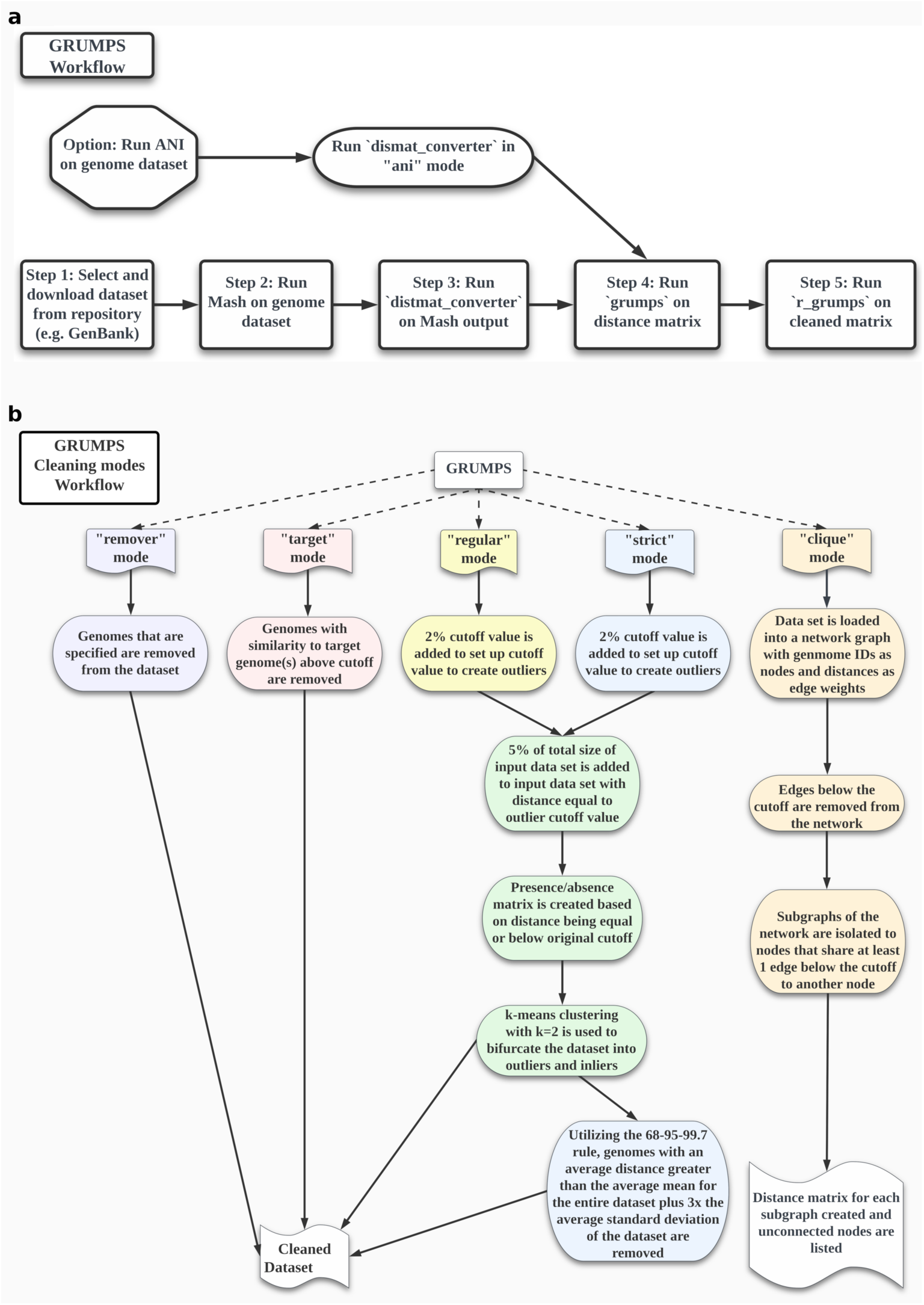
Graphical Summary of GRUMPS workflow and modes. Panel **a** contains a high-level overview of the GRUMPS workflow while panel **b** contains a summary of the major cleaning modes provided by GRUMPS. **a.** Each primary step is indicated by a square and move sequentially from left to right. Optional supported entry points in the workflow are represented by octagonal shapes. Oval shapes indicate the required steps needed to transition from the corresponding optional entry point to the primary GRUMPS workflow. **b.** For each mode, a generalized summary of the steps used in that mode are listed in order. Black arrows indicate the flow from start to finish for each mode. The steps of each mode are colored with a unique color except for the steps shared between “regular” and “strict” modes which are colored green.

### Validation of the Methodology behind GRUMPS

While GRUMPS was developed and tested utilizing a variety of datasets containing multiple bacterial species from different phyla, a collection of three different bacterial species from the phyla Proteobacteria, Bacillota, and Actinomycetota (*Escherichia coli*, *Enterococcus faecium*, and *Mycobacterium abscessus* respectively) were selected for validation. These species were chosen because their population structures have been previously published, allowing for the validation of GRUMPS’ methodology by comparing its results with already established population structures.

### Escherichia coli validation case

A dataset comprising 23,532 *E. coli* genomes was used to demonstrate the utility of both “regular” and “strict” cleaning modes of GRUMPS, using the default cutoff of 0.05. The heatmaps depicted in Figure 3 represent three different states of the dataset. Figure 3a displays a clustered heatmap of Mash pairwise distances among the 23,532 *E. coli* assemblies without being processed by GRUMPS (unclean dataset). Figure 3b illustrates the same dataset after processing with the “regular” mode of GRUMPS and applying the optional “sigma” cleaning step. Figure 3c shows the dataset after employing the “strict” mode of GRUMPS (using the default cutoff) with the “sigma” cleaning step also applied. The values for the mean, maximum, and standard deviation are provided for each mode in the table of Figure 3. After applying the “regular” mode to the initial dataset, 245 genomes were removed, which reduced the mean and maximum Mash pairwise distances by 0.00064 and 0.95897, respectively. After the application of the “strict” mode to the initial dataset, 301 genomes were removed, reducing the mean and maximum Mash pairwise distances by 0.00065 and 0.95989, respectively. The “strict” mode removed an additional 56 genomes compared to the “regular” mode and achieved a mean value 0.00001 and a maximum value 0.00092 smaller than those of “regular” mode. The results obtained by both “regular” and “strict” modes of GRUMPS are consistent with those published by Abram *et al* in 2021^15^ (Supplemental Figure 1a), despite utilizing a dataset twice the size. The heatmap in Supplementary Figure 1b illustrates the distribution of the 23,287 *E. coli* genomes across the previously described 14 phylogroups.

**Figure 3.**
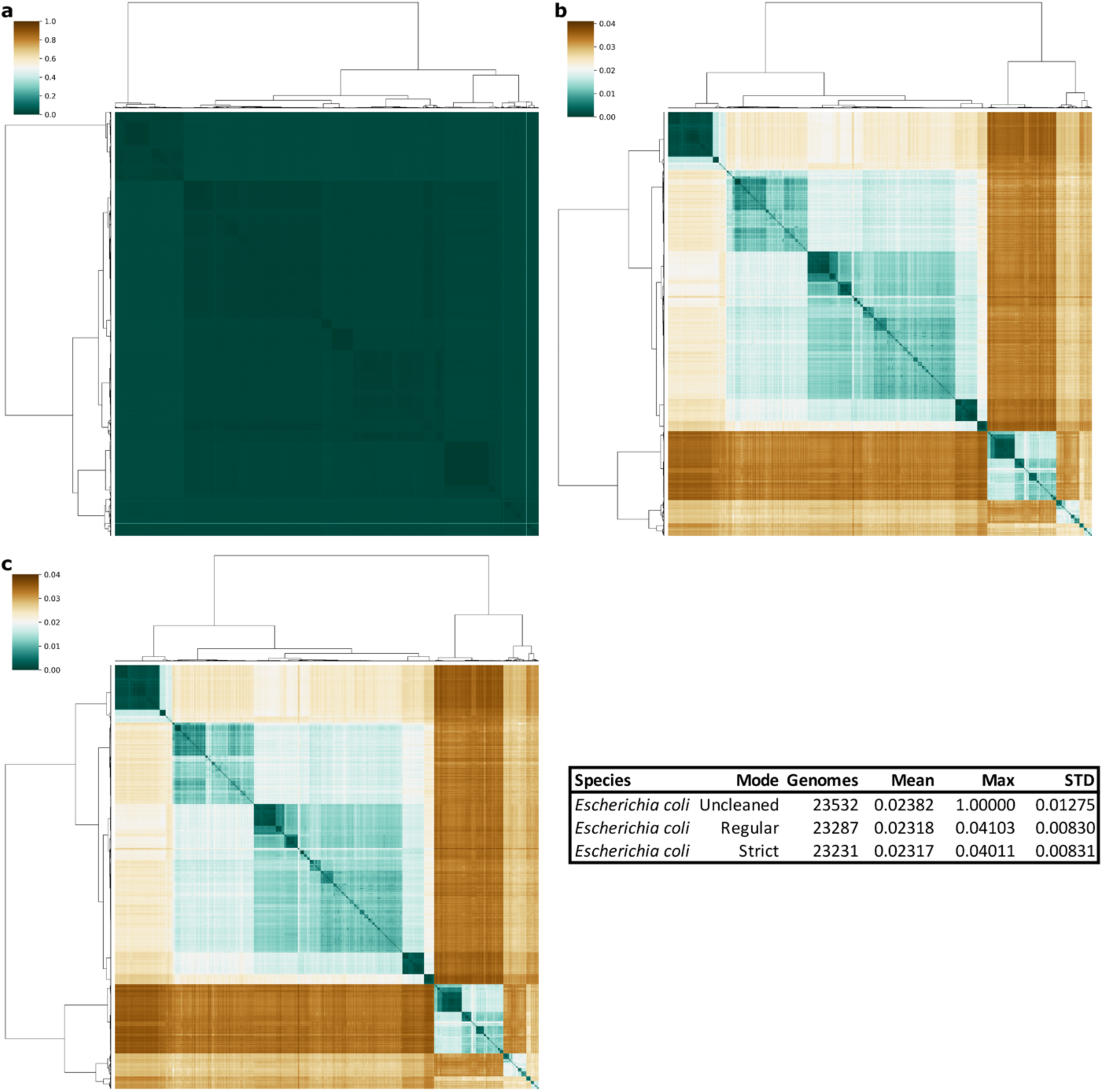
*Escherichia coli* genome datasets before and after cleaning with GRUMPS. Panel **a** displays a heatmap of pairwise Mash distance values for a set of 23,532 *E. coli* genomes. The dataset, not cleaned with GRUMPS, contains many outlier genomes beyond the species cutoff of 0.05. Panel **b** displays a heatmap for a set of 23,287 *E. coli* genomes that has been cleaned using GRUMPS in “regular” mode with the optional “sigma” cleaning step, resulting in the removal of the majority of the outlier genomes. The population structure of the species becomes evident. Panel **c** features a heatmap for a set of 23,231 *E. coli* genomes cleaned using the “strict” mode of GRUMPS, with the optional “sigma” cleaning step. All outlier genomes have been removed from the dataset, showcasing a well-defined population structure of the *E. coli* species, similar to the results obtained using a smaller dataset of 10,667 *E. coli* genomes^15^.

### Enterococcus faecium validation case

To broaden the scope of the validation process, we utilized a dataset of 1,856 *E. faecium* genomes to demonstrate the efficacy of both the “regular” and “strict” cleaning modes of GRUMPS, with the optional “sigma” cleaning step using the default cutoff of 0.05 for Mash distance (Figure 4). In the “regular” mode, 78 genomes were removed, which reduced the mean and maximum values by 0.00307 and 0.21258 respectively. The “strict” mode, further removed an additional 134 genomes, lowering the mean and maximum values by 0.00483 and 0.03264, respectively.

The population structure *E. faecium* is typically divided into three clades: A1, A2, and B. A study^27^ by Udaondo *et al.* (2021) investigated the population structure of *E. faecium* using FastANI and a dataset of 2,273 genomes from GenBank, to classify genomes to these clades. Using these classifications, we compared the described population structure with the results from GRUMPS. Genomes from clade B are localized in the small square in the upper left-hand corner of Figure 4b. Genomes from clades A1 and A2 correspond to the left and right sides, respectively, of the bifurcated branch of the dendrogram of Figure 4b and 4c. The exclusion of clade B genomes when using the “strict” cleaning mode of GRUMPS is attributed to a significant sequencing bias towards genomes belonging to clades A1 and A2 which are genetically more similar, as illustrated by the color variations in the heatmap of Figure 4b. This sequencing bias, along with their closer genetic similarity, shifts the mean Mash distance of the dataset towards clades A1 and A2, leading to the removal of clade B genomes under the three-sigma rule used in the “strict” cleaning mode.

The “regular” cleaning mode of GRUMPS using the default cutoff, aligned best with the currently accepted population structure of the species, distinctly delineating the three known clades (A1, A2 and B). However, it is worth noting that recent genomic studies suggest that strains classified as *E. faecium* clade B should be reclassified as members of the species *Enterococcus lactis* due to their closer genomic proximity to the type strain of *E. lactis* and the absence of hospital-associated markers found in *E. faecium* clade A1 and A2 strains^28–30^. To further investigate the relationships between clade B *E. faecium* strains and *E. lactis*, we utilized two type strain genomes for both species (GenBank Accessions: GCA_001544255.1, GCA_900447735.1, GCA_015904215.1, and GCA_015751045.1) and compared them to all clade B strains. The Mash distances ranged from 0.00947898 to 0.0232342 for comparisons with *E. lactis*, which is noticeably lower than the Mash distances to the type strain genomes of *E. faecium* which had Mash distances to all clade B genomes ranging from 0.040365 to 0.0488843.

**Figure 4.**
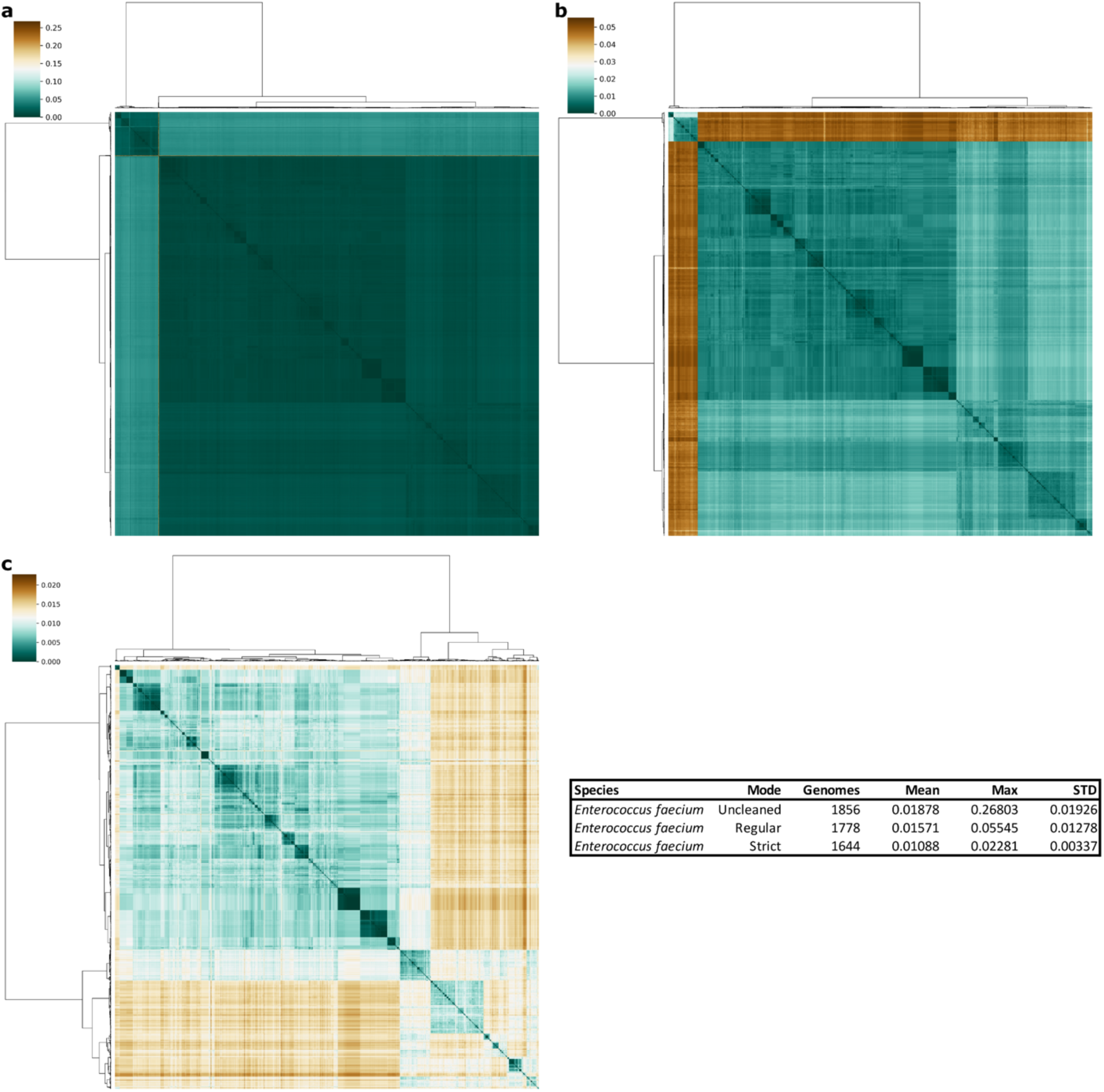
*Enterococcus faecium* genome datasets before and after cleaning with GRUMPS. Panel **a** displays a heatmap for a set of 1,856 *E. faecium* genomes. This dataset has not been processed and includes many outlier genomes, obscuring the population structure. Panel **b** shows a heatmap for a set of 1,778 *E. faecium* genomes. This dataset has undergone cleaning using GRUMPS “regular” mode with the optional “sigma”, resulting in the removal of 78 outlier genomes. The population structure of the species is now more apparent, with clade B distinctly differentiated from clades A1 and A2. Panel **c** features a heatmap for a set of 1,644 *E. faecium* genomes. This dataset has been cleaned using GRUMPS “strict” mode with the optional “sigma” cleaning step which excluded genomes from clade B, enhancing the visibility of the population structure of *E. faecium* clades A1 and A2. The values of the mean and maximum Mash distances, as well as the standard deviation are provided for each mode in the table.

### Mycobacterium abscessus validation case

A set of 1,692 *M. abscessus* genomes was used to illustrate the effectiveness of both “regular” and “strict” cleaning modes of GRUMPS, using the default cutoff of 0.05 and the optional “sigma” cleaning step (Figure 5). After applying the “regular” mode to the initial dataset, 27 genomes were removed, which decreased the mean and maximum values by 0.00031 and 0.03131 respectively. The “strict” mode further reduced the dataset to 1,641 genomes by removing 51 genomes leading to reductions in the mean and maximum values by 0.00041 and 0.03355, respectively.

The results obtained from both, the “regular” and “strict” modes of GRUMPS, distinctly showcase the population structure of *M. abscessus*, with each of its three recognized subspecies (*M. abscessus* subsp. abscessus*, M. abscessus subsp. bolletii and M. abscessus subsp. massiliense)* forming unique cluster within the heatmaps of Figure 5b and 5c. The first square in the upper left-hand corner of the clustered heatmap contains approximately 500 genomes of *M. abscessus* subsp. *massiliense*. The middle square of Figure 5c encompasses around 200 genomes belonging to *M. abscessus* subsp. *bolletii*, while the largest square in the bottom right corner of the clustered heatmaps includes roughly 1,000 genomes of *M. abscessus* subsp. *abscessus*. Subspecies information for these genomes was confirmed by extracting the subspecies metadata for each genome.

Additionally, we leveraged the classification scheme from a study^31^ by Matsumoto *et al.* (2019), which utilized a multilocus sequence type (MLST) based methodology, *mlstverse*, to differentiate these three subspecies from each other. A significant divergence between the results from Matsumoto *et al.* and those obtained by GRUMPS was the absence of an intermixed subpopulation comprising 43 genomes (28 *M. abscessus* subsp. *massiliens* and 15 *M. abscessus* subsp. *abscessus*) that was identified by mlstverse^31^. This discrepancy highlights the higher resolution of whole genome sequencing methods compared to typing methodologies that utilize only a subset of genes, a finding supported by other studies^10,15,27^. Ultimately, GRUMPS provided a clearer classification of *M. abscessus* genomes into their three recognized subspecies (Figure 5).

**Figure 5.**
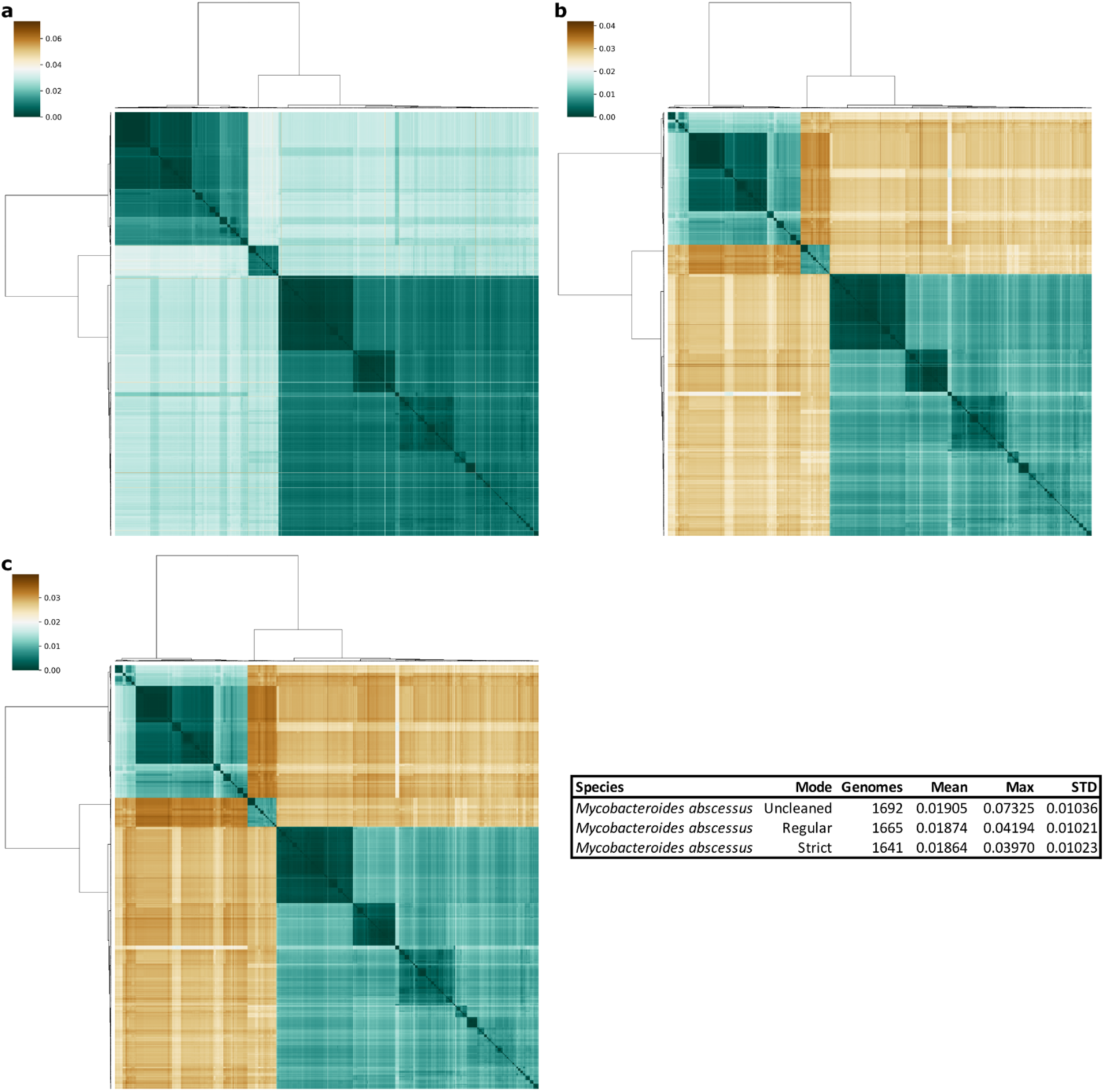
*Mycobacterium abscessus* genome datasets before and after cleaning with GRUMPS. Panel **a** features a heatmap of pairwise Mash distance values for a dataset of 1,692 *M. abscessus* genomes. This dataset has not been processed with GRUMPS and includes genomes that are outside the species cutoff of 0.05. Panel **b** displays a heatmap for a set of 1,665 *M. abscessus* genomes, cleaned using GRUMPS in “regular” mode with the optional “sigma” cleaning step resulting in the removal of 27 outlier genomes. The enhanced clarity in this panel illustrates the underlying population structure more effectively. Panel **c** shows a heatmap for a set of 1,641 *M. abscessus* genomes processed using GRUMPS in “strict” mode with the optional “sigma” cleaning step which removed a total of 51 outlier genomes. This panel offers a clearer visualization of the population structure, with the three subspecies of *M. abscessus* distinctly visible. The values of the mean and maximum Mash distances, as well as the standard deviation are provided for each mode in the table.

### Precision, Recall, and Accuracy Analyses

To evaluate the performance of the two primary cleaning methods offered by GRUMPS, we utilized a dataset comprising 45,558 genomes from *E. coli*, *Salmonella enterica*, and *Staphylococcus aureus.* Then we constructed a series of confusion distance matrices to analyze the precision, recall, and accuracy of both the ‘regular’ and ‘strict’ cleaning modes of GRUMPS. For each of these three species, ten confusion distance matrices were constructed that contained all the genomes of that species in GenBank (as of April 6, 2020). Additionally, each matrix included confusion data consisting of randomly selected genomes from the other two species, where the size of the confusion data represented 10% of the total number of genomes of the species being analyzed, with 5% from each of the other two species.

Each confusion distance matrix was then cleaned using both the ‘regular’ and ‘strict’ cleaning modes of GRUMPS. To determine the accuracy, specificity, and sensitivity of each cleaning mode for each species, the output of GRUMPS for each species dataset and the respective cleaning mode was treated as the ground truth. Accordingly, any genome present in both, the output of GRUMPS for the species and the confusion datasets for that species, was considered a true positive. Genomes that appeared only in the output from the confusion distance matrix were classified as false positives. Remarkably, for both cleaning methods across all three species, the measures of accuracy, sensitivity, and specificity reached 100%. Given that all 30 confusion distance matrices consistently yielded the same high levels of accuracy, sensitivity, and specificity for both cleaning methods, we conclude that the primary cleaning methods provided by GRUMPS are both highly reliable and replicable.

### Validation of Sigma Mode

To demonstrate the impact of the optional “sigma” cleaning step available in both “regular” and “strict” cleaning modes of GRUMPS, we applied the “regular” cleaning mode to an *E. coli* dataset with and without the optional “sigma” cleaning step. The resultant heatmaps are presented in Figure 6 with panel 6a displaying the heatmap generated by “regular” mode without “sigma” cleaning and panel 6b showing the heatmap generated by “regular” mode with “sigma” cleaning. The population structure of *E. coli* is more distinctly illustrated in the clustered heatmaps when “sigma” cleaning is employed, despite this step removing only an additional 0.8% genomes. The color scale of the cluster heatmap, which is automatically generated based on the minimum and maximum values in the dataset, becomes more effective once genomes that deviate from the expected distribution of values for the species are removed. This removal enhances the clarity of the population structure within the heatmap.

The distribution of pairwise comparisons for both datasets is shown in Figures 6c and 6d respectively. A comparison of the topology of the histogram for both datasets, reveals that the “sigma” cleaning step effectively removes outlier genomes without altering the overall population structure, which is represented by peaks and valleys in the histogram. Additionally, the “sigma” cleaning step reduces the maximum value for the *E. coli* dataset to 0.04103, which is below the species boundary of 0.05, whereas the “regular” cleaning mode without the “sigma” step results in a maximum value of 0.07234, exceeding the species boundary.

**Figure 6.**
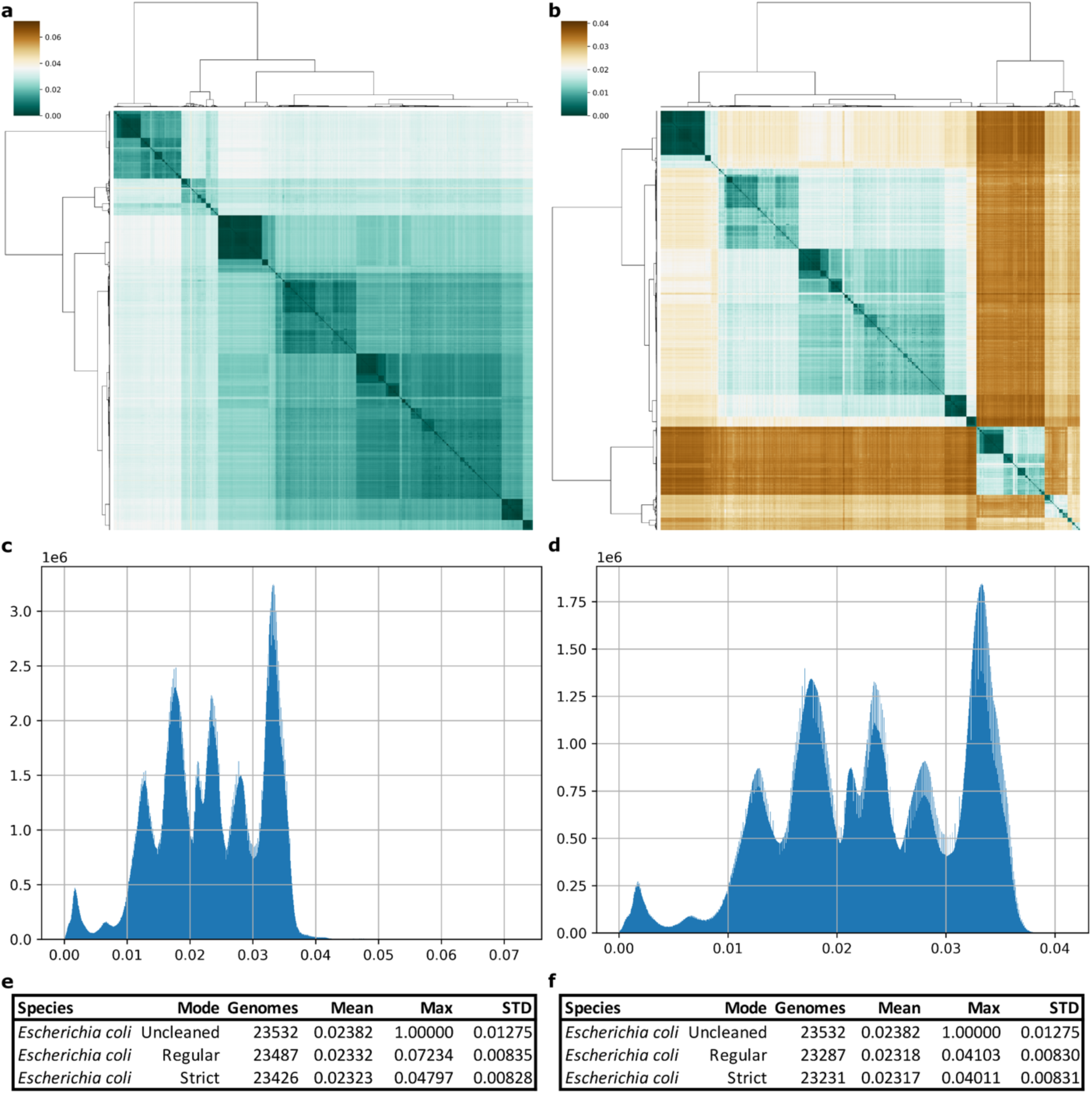
Comparison of GRUMPS “regular” cleaning mode with and without “sigma” filtering using *Escherichia coli*. Panel **a** displays a heatmap of pairwise Mash distance values for a dataset of 23,487 *E. coli* genomes. This dataset was processed using the “regular” mode of GRUMPS without “sigma” cleaning, including pairwise comparisons that exceed the species boundary of 0.05. Panel **b** features a heatmap generated by using the “regular” cleaning mode of GRUMPS with “sigma” cleaning, showing no pairwise comparisons above the species boundary, thereby enhancing clarity. Panels **c** and **d** show histograms of the pairwise comparisons used to produce panels **a** and **b** respectively. Panel **e** contains the statistical data: number of genomes, mean, maximum value, and standard deviation, for the initial (uncleaned) dataset, with “regular” mode, and “strict” mode without “sigma” cleaning. Panel **f** provides the same statistical information with “sigma” cleaning.

### Effect of Plasmid Sequences on GRUMPS

While GRUMPS provides a rapid methodology for cleaning datasets, it is important to note that the outcome is primarily influenced by the method (ANI or Mash distance) utilized to determine the genetic similarity between a pair of genomes. This in turn defines the impact of including or excluding plasmid sequences in the output from GRUMPS. Utilizing statistical cleaning methods and unsupervised machine learning based on these similarity values, GRUMPS decides on the inclusion or exclusion of genomes from its output.

To investigate the impact of plasmid sequences on our methodology, we developed a dataset of 877 complete *E. coli* genomes, that included plasmids, and created a duplicate set of genomic sequences that had all plasmid sequences removed. After applying both ‘regular’ and ‘strict’ cleaning modes of GRUMPS, the removal of plasmid sequences led to a 0.228% variation between the two datasets. Specifically, two genomes (GCA_900088105.2 and GCA_900184875.1) were present in the dataset with plasmid included but absent in the dataset without plasmids after applying the cleaning. This suggests that excluding plasmid sequences results in a minor variation in the output, with minimal effect as the size of the dataset increases. However, it should be emphasized that the sensitivity to a plasmid inclusion or exclusion ultimately depends on the tool used to obtain the genetic similarities provided, which is outside the scope of GRUMPS.

### Computational Performance of GRUMPS

To benchmark the computational efficiency of GRUMPS’ in its “summary”, “regular”, and “strict” modes, we utilized a series of distance matrices comprising of *E. coli* genomes, ranging in size from 500 to 23,532, with increments of 500 genomes. Performance testing for both “regular” and “strict” cleaning modes, was conducted with and without the optional features available for these modes (see Methods). The wall time for “summary” mode varied from 2.7 to 332.9 seconds. For the “regular” mode without the heatmap and sigma options, the wall time ranged from 2.4 to 1081.7 seconds, while the “strict” mode without these options showed similar times (from 2.4 to 1082.4 seconds). Including the heatmap and “sigma” options affected the wall times, with the heatmap option increasing times significantly, from approximately 2 to 424 seconds depending on the dataset size. The “sigma” option had a minimal impact with times ranging from 2.4 to 1083.9 seconds. Processing the largest dataset of 23,533 *E. coli* genomes with both, heatmap and sigma options enabled, took approximately 25 minutes. These benchmarks were performed on a workstation with the following configuration: Ubuntu 22.04.1, 2.1 GHz Intel Xeon Gold 6230, 200GB of Micron DDR4 2933 RAM, and a 1TB Micron 2200S NVMe.

## Conclusions

GRUMPS represents a robust data cleaning platform that utilizes statistical analyses and machine leaning to identify and remove low-quality and misidentified genomes. This novel methodology allows for the rapid and reproduceable cleaning of bacterial genomic datasets of any size. The precision, recall, and accuracy analyses, along with its comparable performance on real world datasets to previously published results, underscore the reliability of the methodology.

To evaluate GRUMPS’ capability to accurately identify and remove low-quality genomes from species level datasets, several validations analyses were performed. We selected three bacterial species belonging to three unique phyla, Proteobacteria, Bacillota, and Actinomycetota, to assess GRUMPS’ efficacy on real world datasets.

Initially, the results from GRUMPS’ “regular” cleaning mode were compared with the results obtained in the largest *E. coli* analysis conducted to date^15^ using a dataset twice the size. GRUMPS successfully reproduced the distribution of *E. coli* genomes into 14 previously described phylogroups, as demonstrated in Abram et al., 2021 (Figure 3 and Supplementary Figure 1).

Further validation was pursued by comparing GRUMPS results for both *E. faecium* and *M. abscessus* against recent studies^27,31^ that explored the population structure of these two species. For *M. abscessus*, both the “regular” and “strict” cleaning modes revealed the known population structure, distinctly separating the recognized subspecies. In the case of *E. faecium*, the “regular” cleaning mode disclosed the three recognized clades of the species^27,32^. However, the “strict” cleaning mode removed all clade B strains from the dataset. Although this might seem like overcleaning, recent genomic studies on clade B strains of *E. faecium* suggest that these strains should be reclassified as members of the species *E. lactis*, a position supported by our findings on the genetic similarity between clade B strains and type strains for both *E. faecium* and *E. lactis*^28^.

Benchmarking results demonstrated that GRUMPS is a computationally efficient, capable of processing over 23,000 *E. coli* genomes in less than 30 minutes using local computational resources. The primary computational bottlenecks are the size of the input data, which affects read and write time and overall wall time of the cleaning process, and the disk speed of the storage device used for the dataset.

In a nutshell, GRUMPS is impactful for three main reasons: its ability to accurately identify and remove low-quality genomes from species-level datasets, isolate species-level datasets from genus level datasets, and rapidly elucidate the population structure of any bacterial species. This methodology will greatly aid researchers in obtaining high-quality datasets for comparative genomics studies, even when handling large-scale data. Additionally, GRUMPS and its helper scripts provide an easy-to-use platform for generating high-quality figures and exploring the population structure of any bacterial species. Ultimately, the advanced capabilities of GRUMPS will not only streamline large-scale genomic analyses, providing researchers with highly curated datasets, but also pave the way for revolutionary discoveries in the area of microbiology. For instance, GRUMPS can be used to discover new bacterial species -genomospecies, or species that can be differentiated from other species using genotypic means, specifically genomic methods using genus-level noisy data, while adhering to the established genomic thresholds such as 0.95 ANI^12,22^. Moreover, the analysis of bacterial population structures will allow tracing of bacterial evolutionary histories, revealing how different strains or subspecies have adapted to their environments or hosts. This understanding is vital to elucidate the roles and interactions of bacterial groups within ecosystems, highlighting their contributions to or disruptions of ecological balances. Additionally, the study of population structures is crucial for cataloging biodiversity, contributing significantly to our understanding of global microbial biodiversity. Therefore, the results provided by GRUMPS will have potential applications in diverse areas including medicine, industry, and agriculture.

## Methods

### Test datasets

To conduct the analysis, the prokaryotes summary table was downloaded from NCBI on April 6, 2020; the total number of entries in GenBank on this date was 248,003. The table was then filtered to only include entries with ‘Bacteria’ in the ‘Organism Groups’ column resulting in a set of 243,260 possible genomes. To reduce the impact of low-quality genomes, we exclude genomes with more than 500 contigs which resulted in a reduced dataset of 231,608 genomes. To reduce the impact of uncultured genomes, we removed any genome with ‘uncultured’ in the ’#Organism Name’ column which resulted in a reduced final set of 218,099 potential genomes.

To generate an approximate abundance of species within this reduced dataset, the ’#Organism Name’ column was isolated and each entry was split by white space and only entries with one or more white spaces were retained. Next, we joined the first two splits with a single white space and appended them to a new list. We then trimmed any leading or trailing ‘ characters to improve the accuracy of counting unique genus/species strings and counted the frequency of each unique genus/species string; the number of unique genus/species strings was 13,802. To reduce the impact of poorly annotated genomes, we removed any genus/species strings that contained either ‘bacterium’ or ‘sp.’ which resulted in a set of 11,913 unique genus/species strings.

In order to demonstrate the scalability of GRUMPS, we assessed the population structure of at least 50% of the final reduced set of bacterial genomes by including the most numerous species until there were at least 109,050 genomes in the test set. This resulted in a test set containing 31 species of bacteria and 110,220 genomes. It should be noted that based on the results of our previous work and in accordance with the literature, we considered genomes belonging to the genus *Shigella* to be within the species *Escherichia coli* and counted all *Shigella* labeled genomes and *E. coli* labeled genomes as the same species. We then used the GCAs of all 110,220 genomes in the test dataset to download the genomic .fna file using BatchEntrez.

### Plasmid Dataset

Using the metadata provided by NCBI, we identified a set of 877 complete *E. coli* genomes that contained plasmid sequences and we created a duplicate copy for each genomic file that excluded all plasmid sequences. Mash sketches (k=21 and s=10,000) were generated using all 877 genomes: a set that included the plasmid sequences and a set with the plasmid sequences removed. Mash distances were obtained for both sets and converted into a distance matrix using the helper script provided in GRUMPS. We then utilized both ‘regular’ and ‘strict’ cleaning modes of GRUMPS on each dataset. Both cleaning modes were compared by identifying any genomes that were present in the plasmid included dataset but missing from the plasmid removed dataset.

### Precision, Recall, and Accuracy Analyses

We identified the three most numerous species (*E. coli, Salmonella enterica,* and *Staphylococcus aureus*) in our test data to create multiple sets of confusion data to assess the accuracy, sensitivity, and specificity of GRUMPS. Using the sketches generated in the initial test datasets, pairwise Mash distances for all 45,558 genomes in these three species were obtained. These distances were then converted into a distance matrix using the provided helper script. To create 10 confusion datasets for each species, we randomly selected genomes totaling 10% of the total size of the species under investigation from the other two species, 5% from each. This resulted in a set of 30 confusion distance matrices that were then cleaned using both ‘regular’ and ‘strict’ cleaning modes implemented in GRUMPS resulting in 60 total sets. To calculate accuracy, specificity, and sensitivity of both cleaning modes per species, we considered the output of each cleaning method from the original test data of that species to be the ground truth, meaning any genome retained by the respective cleaning method in the original test data analysis were considered true positives and any genome removed by the respective cleaning method in the original test data analysis were considered true negatives. For each of the 60 sets, true positives were genomes that were present in both the original set and the test set, true negatives were genomes that were absent in both the original set and the test set, false positives were genomes that were present in the test set but absent in the original set, and false negatives were genomes that were removed in the test set but present in the original set.

### Benchmarking

To benchmark the speed of GRUMPS’ “summary”, “regular”, and “strict” modes, we produced a set of distance matrices with genomes ranging from 500 to 23,532 with a step size of 500 from the *E. coli* dataset used in the validation section. For each mode, we ran GRUMPS 10 times and recorded the wallclock time which was then averaged to obtain the wallclock time for each cleaning mode. For both “regular” and “strict” cleaning modes, we utilized the following different sets of optional features to assess the computational increase associated with the optional features: with heatmap with sigma, with heatmap without sigma, without heatmap with sigma, and without heatmap without sigma. Benchmarking was performed using a workstation with the following configuration: Ubuntu 22.04.1, 2.1 GHz Intel Xeon Gold 6230, 200GB of Micron DDR4 2933 RAM, and a 1TB Micron 2200S NVMe. A MacBook Pro (Mid 2015 model) with 16GB of 1600MHz DDR RAM, a 2.8GHz Quad-Core Intel Core i7 processor (i7-4980HQ), and a 1TB Apple SATA SSD SM1024G took approximately 55 minutes to run GRUMPS in ‘regular’ mode with both the heatmap and sigma filtering options enabled on the full *E. coli* dataset containing 23,532 genomes.

### GRUMPS Workflow

An overview of the typical GRUMPS workflow is shown in Figure 2a. In summary, researchers need to identify and download the genomes they intend to analyze before users run Mash (or another comparison method which has definitive species-level cutoffs such as ANI) on the genome set. Based on extensive testing, we generally recommend that genomes are sketched first using a k-mer size of 21 and 10,000 sketches before Mash distances are calculated to increase both performance and reproducibility; however, it is possible that these settings will not work for every bacterial genome that has or will be sequenced and different settings might be required depending on the dataset of interest. Once Mash distances are obtained, the provided helper script ‘distmat_converter’ can be utilized to create a NxN symmetrical distance matrix (where N is the number of genomes) required by ‘grumps’. If a comparison method other than Mash is utilized, such as ANI, the helper script ‘distmat_converter’ should be run with ‘-m ani’. When ‘distmat_converter’ is run with ‘-m ani’, users can either convert their values to Mash distances with ‘-c yes’ or invert their values with ‘-i yes’. Only one of these options can be used at a time and ‘-c yes’ is prioritized over ‘-i yes’. Converting to Mash distances is accomplished by the following: 1-(ANI/100). Inverting is accomplished in a similar fashion: 100-ANI. We highly recommend users to convert their values to Mash distances as GRUMPS was designed to work with values in this range. If users choose to invert instead of convert, users will need to manually specify their cutoff to an appropriate value. A cutoff of 5 would correspond to a species boundary of 95% ANI value. Once users have a distance matrix, one of the available cleaning methods described in detail below can be leveraged to clean the dataset. Finally, the supplied R script ‘r_grumps’ can be used to produce graphical representations of the dataset. The available cleaning modes in GRUMPS and their function are summarized in Figure 2b.

## Author contributions

Conceptualization: K.Z.A.

Data curation: K.Z.A.

Formal Analysis: K.Z.A.

Funding acquisition: S.J.

Investigation: K.Z.A. Z.U.

Methodology: K.Z.A.

Software: K.Z.A.

Supervision: S.J. Z.U.

Validation: K.Z.A. Z.U.

Visualization: K.Z.A. Z.U.

Writing-original draft: K.Z.A. Z.U.

Writing – review & editing: K.Z.A. Z.U. M.R. S.J.

## Competing interesting

Authors declare no competing interests.

## Acknowledgments

A portion of the analyses were performed with the GRACE cluster, provided by the University of Arkansas for Medical Sciences (UAMS) and managed by the Department of Biomedical Informatics. We are thankful to Intawat Nookaew for discussions and suggestions provided to early versions of this manuscript.

## Supplementary Information

**Figure S1.**
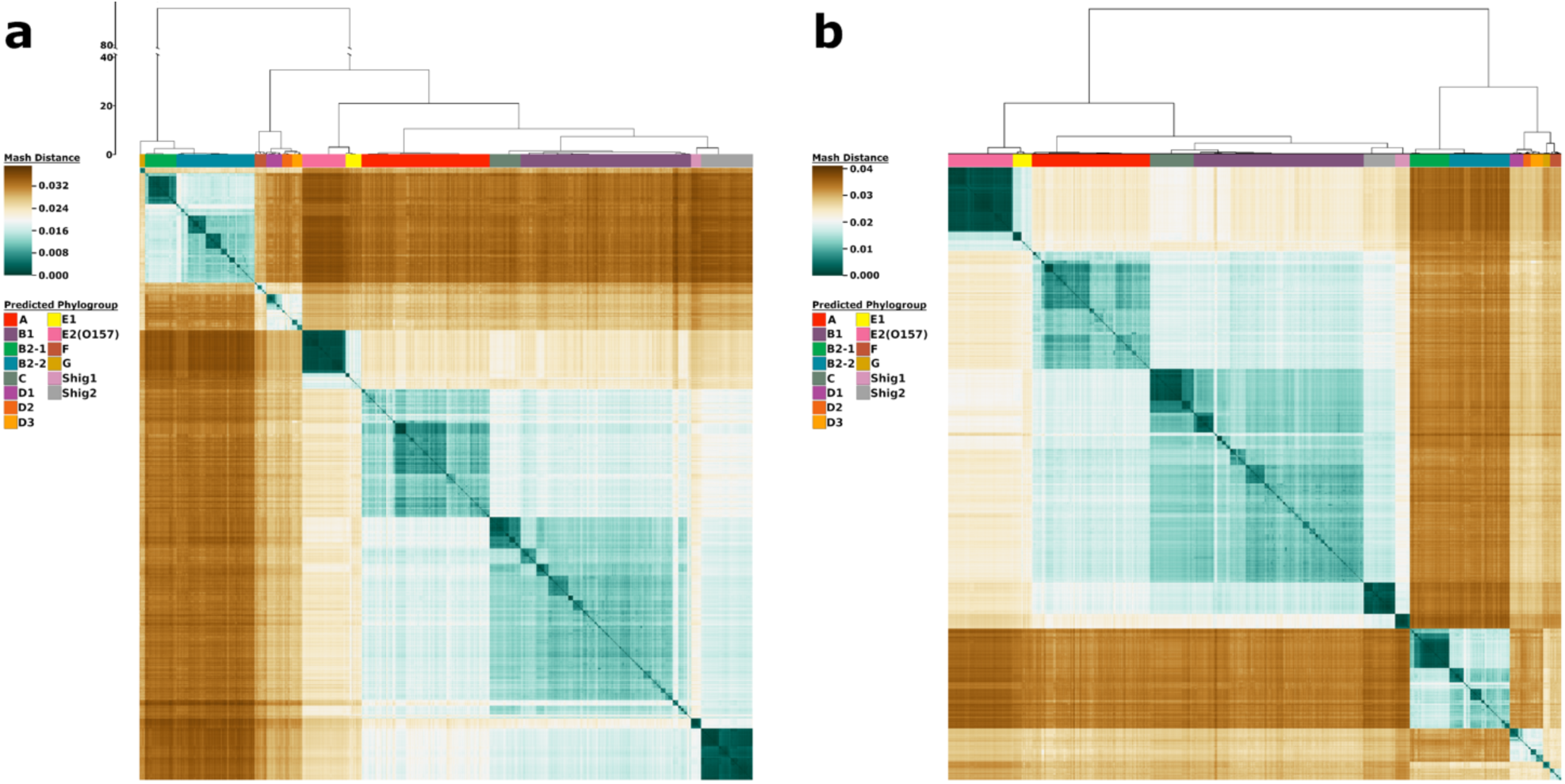
Comparison of *Escherichia coli* population structure from Abram et al. 2021 and GRUMPS “regular” mode. Panel **a** contains a heatmap of pairwise Mash distance values for a set of 10,667 *E. coli* genomes published in Abram et al. 2021. Panel **b** contains a heatmap of pairwise Mash distance values for a set of 23,287 *E. coli* genomes produced by GRUMPS with “regular” cleaning mode and “sigma” filtering. The color bars at the top of both heatmaps identify the phylogroups as predicted in Abram et al. 2021. The colors in the heatmap are based on the pairwise Mash distances. Shades of teal represent similarity between genomes, with the darkest teal corresponding to identical genomes reporting a Mash distance of 0. Shades of brown represent low genetic similarity per Mash distance, with the darkest brown indicating a maximum distance of ∼ 0.039. Genomes of relative median genetic similarity have the lightest color. While the ordering of the phylogroups differs between panel **a** and **b**, the classification remains the same.

