## Supplementary Material for "Genomic Distance-based Rapid Uncovering of Microbial Population Structures (GRUMPS): a reference free genomic data cleaning methodology"

**Detailed description of the different modes available in GRUMPS.** The “summary” mode is the intended starting point for any cleaning effort leveraging GRUMPS. This mode provides multiple statistical summaries of the input dataset as well as a histogram of the comparison values in the dataset and enables the manual identification of outlier genomes, if desired. Based on internal testing, if a cutoff value different from the default value of 0.05 is desired, we recommend using the sum of the mean and standard deviation output by the “summary” mode of GRUMPS as the cutoff value for a given dataset. This can be useful when analyzing species, such as *M, tuberculosis,* whose genomic variability has been documented to be relatively smaller than other species^26^ or datasets with enough contamination that the overall average of the dataset is greater than 0.05.

The “regular” and “strict” modes are the primary cleaning methodologies provided by GRUMPS. Both modes function by adding a small number of artificial outlier comparisons (5% of the total input dataset size) to the dataset where the pairwise values of all artificial outlier comparisons to all initial members of the dataset to be cleaned, are set to slightly above the cutoff (102% of the specified cutoff value). This creates a very small set of controlled artificial outliers that are just outside the boundary defined by the cutoff value for the species level, thus increasing the utility and reliability of the subsequent clustering step. Once the artificial outliers are added, both, "regular" and "strict" cleaning modes utilize k-means clustering (k=2) to produce two clusters, one cluster containing inlier genomes and the remaining cluster containing outlier genomes as well as the artificial outliers inserted by GRUMPS. The "strict" cleaning mode of GRUMPS then removes any genome whose mean distance to all other genomes in the dataset is greater than the sum of the mean and three times the standard deviation (three-sigma rule). For both “regular” and “strict” cleaning methods, an additional optional cleaning step, described below, can be applied.

The “sigma” mode is an alternative cleaning method which leverages the three-sigma rule on the distribution of comparisons for each genome to remove genomes with divergent behavior on either side of their distribution of comparisons relative to the rest of the genomes in the dataset. In this cleaning step, the distribution of comparison values for each genome is isolated and the extreme left and right (defined as 0.03% on the left and 99.7% on the right) tails of each distribution are identified and a cutoff for each side is obtained by summing the mean of that side plus three times the standard deviation. By default, this cleaning mode is automatically applied as an additional cleaning step at the end of the “regular”, “strict”, “target”, and “remover” cleaning modes.

The “target” mode is intended to be used with one or more reference genomes (such as type strain assemblies or other genomes of interest) and removes any genomes from the dataset that do not have a distance value less than or equal to the set cutoff to all the reference genomes supplied. The identifier for each reference genomes utilized for “target” mode must be in columns and rows of the distance matrix that is the input to GRUMPS.

The “remover” mode is intended to remove one or more specified genomes from the input dataset which might consist of either easily identified or known outlier genomes. These genomes can be manually identified using the output of “summary” mode or if the researcher has conducted additional analyses indicating that a genome in their initial dataset should not be included in the cleaned dataset.

The “clique” mode is the final primary mode provided by GRUMPS and uses a graph-based clustering approach to clean datasets. This mode is intended to be used on datasets that contain multiple species (such as an entire genus). The “clique” mode partitions the dataset into multiple smaller uncleaned species level datasets as well as a set of unconnected genomes. The initial dataset is converted into a graph where nodes are genomes and edges are weighted by the distance between pairs of genomes. Utilizing the specified cutoff, GRUMPS trims all edges that are greater than the cutoff and returns all subgraphs. The distance matrix for each subgraph (presumably, species) identified by this mode is output, as well as a list of genome sequences that have no connections to any genome in the dataset. These distance matrices should be considered uncleaned and should be fed back into GRUMPS to be cleaned by one of the two primary cleaning modes of GRUMPS, “regular” or “strict”, before subsequent analyses are performed on these datasets.

When utilizing “clique” mode, it is possible that some resultant datasets are not large enough for “regular” or “strict” mode to partition using the k-means clustering approach implemented in these modes. This can occur when the “clique” mode returns a resultant dataset with a small number of members where 5% of the resultant dataset is less than 1 (for example a resultant dataset with less than 100 members). To address this issue, a “small” cleaning mode is provided by GRUMPS which uses the average distance of each genome (ignoring self-comparisons) to remove outlier genomes. For example, if a dataset containing 95 genomes has 5 genomes with an average distance above the specified cutoff, then those 5 genomes would be considered outliers and removed from the dataset.

An additional optional cleaning step, “medoid” (-M yes) is available for the “regular”, “strict”, and “clique” cleaning modes. For “regular” and “strict” modes, this cleaning step identifies the medoid of the dataset and removes any genomes that have a Mash distance greater than the cutoff (0.05 default) to the medoid of the dataset. For “clique” mode, the “medoid” cleaning step identifies the optimal number of subgroups using K-means clustering and silhouette scores (k ranges from 2 to 10 groups). For “clique” mode, a set of artificial outliers is added in the same manner as for “regular” or “strict” modes. This is done to ensure there are at least 2 subgroups within a given set of connected genomes. Once the optimal number of subgroups are identified, the medoid of each subgroup is identified and subgroups are merged if the medoids of a given subgroup is under the cutoff for species membership (default 0.05).

A high-level overview of the intended GRUMPS workflow is provided in Figure 2a, including an alternative entry point to the workflow which have helper scripts provided to aid researchers in alternate workflows leveraging GRUMPS. A detailed summary of the steps used in each cleaning method provided by GRUMPS is shown in Figure 2b.
